# IL-13 Modulates Antiviral Effector and Proinflammatory Pathways in Rhinovirus-Infected Pediatric Bronchial Epithelium

**DOI:** 10.64898/2025.12.06.692776

**Authors:** Patricia C. dela Cruz, Basilin Benson, Naresh Doni Jayavelu, Weston T. Powell, Lucille M. Rich, Elizabeth R. Vanderwall, Camile R. Gates, Maria P. White, Nyssa B. Samanas, Kourtnie Whitfield, Teal S. Hallstrand, Steven F. Ziegler, Gail H. Deutsch, Matthew C. Altman, Jason S. Debley

## Abstract

**Background:** Rhinovirus (RV) is the most common trigger of viral-induced pediatric asthma exacerbations. The impact of IL-13-driven inflammation, common in pediatric asthma, on airway epithelial antiviral and inflammatory responses to RV remain unclear.

**Objective:** Determine how IL-13-driven T2 inflammation modulates pediatric bronchial epithelial cell responses to RV infection.

**Methods:** Bronchial epithelial cells (BECs) were collected from children with (n=50) and without (n=11) asthma. They were differentiated at an air-liquid interface for 21 days, pretreated with IL13 (10ng/mL) for 7 days to model T2 inflammation, then infected with RV-A16 (MOI 0.5). RNA sequencing of BECs was performed prior to and on days 2, 4, 7, and 10 post infection. Linear and generalized additive models partnered with pathway analysis identified differentially expressed gene clusters.

**Results:** RV infection, IL-13 stimulation, and their interaction each induced differentially expressed genes (7,808; 10,251; and 7,095 genes, respectively; FDR<0.05). IL-13 pretreatment did not alter RV load or a cluster enriched for interferon response and regulation genes, including *IFNB1, IFNL1-3, STAT1/2,* and *CXCL10/11*. In contrast, IL-13 reduced expression of distinct antiviral effectors (e.g. *MX1/2, RSAD2, IFITM1-3*; FDR=1.85×10⁻⁴) and increased a secondary proinflammatory response cluster enriched for IL-15, TNF, ER stress, and cell-death pathways (FDR=3.63×10⁻⁶). These clusters correlated with viral load in RV-infected cells, but IL13 pretreatment eliminated those associations.

**Conclusions:** IL-13 does not modify viral load or interferon induction but selectively suppresses epithelial antiviral effector programs and enhances secondary inflammatory pathways during RV infection. These findings provide mechanistic insight into how T2 inflammation contributes to viral-triggered asthma morbidity.

**Key Messages:**

– IL-13 did not alter rhinovirus viral load or epithelial interferon induction but selectively suppressed antiviral effector programs. This indicates that T2 inflammation reshapes antiviral defenses downstream of interferon signaling rather than impairing interferon production itself.
– IL-13 enhances a distinct secondary proinflammatory cluster enriched for IL-15, TNF signaling, ER stress, and cell-death pathways. These late-phase responses may contribute to the heightened airway inflammation and injury observed during viral-triggered exacerbations in T2-high pediatric asthma.
– IL-13-driven changes decouple epithelial immune responses from viral load, revealing mechanisms that may persist even with anti-IL-4Rα/IL-13 therapy. These findings identify epithelial antiviral effector pathways and IL-15/TNF-associated inflammatory programs as potential therapeutic targets for patients who experience breakthrough viral exacerbations despite IL-13 blockade.

**Capsule Summary:** IL-13 did not alter rhinovirus load or epithelial interferon but selectively suppressed antiviral effectors and amplified secondary inflammatory pathways, revealing how T2-inflammation may worsen viral-triggered asthma exacerbations and contribute to breakthrough exacerbations despite IL-4Rα/IL-13-targeted therapy.

## INTRODUCTION

Asthma continues to be a major medical and economic burden in the United States, affecting 24.9 million people and costing over $80 billion annually (1). Viral infections, most commonly rhinovirus (RV), trigger up to 90% of pediatric asthma exacerbations (2,3). A large portion of pediatric asthma is Type-2 (T2) asthma (4), which is characterized in part by IL-13 driven airway inflammation (4,5). Within the asthmatic airway epithelium, IL-13 exposure has been shown to induce goblet cell hyperplasia and impact epithelial remodeling (4,6,7). Although there is evidence that T2 inflammation may dampen antiviral responses (8), the impact of IL-13 driven inflammation on innate immune responses to RV by the airway epithelium remains poorly understood.

In this study, we assess the impact of T2 inflammation on bronchial epithelial responses and viral load following rhinovirus infection. Prior investigations of the impact of IL-4 and IL-13 on RV replication and epithelial innate immune responses to RV are conflicting, with some studies reporting in primary human bronchial epithelial cells (BECs) that IL-4 or IL-13 conditioning increased RV replication while reducing RV-induced interferon responses, downregulating TLR3, impairing IRF3 activation (9), or increasing epithelial ICAM-1 (the major RV receptor) expression (10). Other data suggest that IL-13 exposure may increase RV-induced CXCL10, IL8, and GM-CSF expression without altering RV replication (11), or that IL-13 reduces RV replication in the setting of induced goblet cell metaplasia, impaired ciliogenesis, and enhanced IFN and ISG responses (12).

Prior studies, including those from our group, have implicated dysregulated interferon responses in viral-induced asthma exacerbations and remodeling responses associated with airway obstruction (8,13–18). Deficient type I and III IFN responses to rhinovirus infection in asthmatic airway epithelium have been reported (13–15). However, this concept is controversial as other groups have not observed deficient IFN responses to viruses (rhinovirus, RSV, influenza) by asthmatic airway epithelium, and indeed have described greater expression of type I and III interferons and associated interferon-stimulated genes following viral infection that correlated with reduced lung function and greater obstruction (16,17,19). Seemingly discordant observations might be explained by differences between studies in the model systems used, clinical characteristics of donors and/or molecular asthma endotypes, and heterogeneity in the kinetics of airway epithelial IFN I/III response between individuals. Two recent corroborating studies showed that in nasal epithelium (18) and bronchial epithelium, specifically non-ciliated cells (20), low interferon expression in the uninfected state was associated with higher rhinovirus replication 2-3d post infection and was associated with more frequent pediatric asthma exacerbation.

In this study, we investigate the link between IL-13 and epithelial interferon responses to viruses in pediatric asthma. Specifically, we hypothesized that IL-13 mediated inflammation alters epithelial responses to rhinovirus, specifically dampening epithelial interferon responses, leading to increased viral load and subsequent enhanced inflammatory and remodeling responses. To test this hypothesis, we utilized organotypic air-liquid interface (ALI) cultures from primary BECs from well characterized children with and without asthma (6,16,21,22) and modeled T2 inflammation by conditioning cultures with IL-13 (23,24). Cultures were subsequently infected with rhinovirus and transcriptomic analysis of epithelial RNA was performed over 10 days following infection to allow for a comprehensive assessment of the kinetics of viral replication and host bronchial epithelial host responses with or without IL-13 conditioning. Our central aim was to elucidate mechanisms by which IL-13 affects epithelial responses to rhinovirus infection to provide novel insights into why rhinovirus infection is associated with severe exacerbation in children with uncontrolled T2-asthma.

## MATERIALS AND METHODS

### Study population & clinical characterization

Experiments were completed under studies #12490 and #1596 approved by the Institutional Review Board at Seattle Children’s Hospital. Assent was given by children over 7 years old and parents of children provided written consent. All pediatric subjects were clinically characterized during an outpatient phenotyping visit. A detailed review of pertinent medical history including history of atopy was performed (asthma exacerbation history and medications, prior allergy testing by skin prick or serum IgE, atopic dermatitis, allergic rhinitis). Criteria for an asthma exacerbation were an increase in asthma symptoms requiring hospitalization/emergency department visit or systemic corticosteroids for three or more consecutive days (25). Spirometry and measurement of fraction of exhaled nitric oxide (FeNO) were performed in accordance with the American Thoracic Society/European Respiratory Society guidelines (26,27), with percent predicted spirometric measurements calculated using race-neutral reference equations defined by the Global Lung Function Initiative (28). Total systemic eosinophil count, serum IgE, and allergen-specific IgE to cat, dog, D. farinae, D. pteronyssinus, aspergillus fumigatus, and timothy grass were quantified in the blood.

### Air-liquid interface (ALI) cultures of bronchial epithelial cells

Bronchial epithelial cells (BECs) were obtained from children with asthma (n=50) and healthy children (n=11) under general anesthesia for elective procedures. 4-mm Harrell unsheathed bronchoscope cytology brushes (CONMED®) were placed through the endotracheal tube as previously described (29). Cells were seeded then proliferated in submerged cultures in collagen Type 1-precoated T-25 culture flasks (Corning®). Following proliferation using PneumaCult™EX-Plus medium (Stemcell™) for 4-7 days, BECs were differentiated to an organotypic pseudostratified ciliated state for 21 days at ALI using PneumaCult™ALI medium (Stemcell™) (16,30,31). Light microscopy of live cell cultures, H&E and immunofluorescent staining, as well as transmission and scanning electron microscopy were performed to assess cell morphology, presence of cilia, and mucous production (See supplemental methods for additional detail).

### IL-13 treatment

Recombinant human IL-13 (10ng/mL, R&D Systems®) was added to the basolateral compartment of differentiated BEC cultures with each media change starting 7 days prior to rhinovirus infection and continued through the completion of all experimental harvest timepoints.

### Human rhinovirus A16 (RV-A16) infection of BECs

RV-A16 (MOI 0.5) was added to the apical surface of BECs in ALI, incubated for two hours, then removed without subsequent rinsing. Genesig® Human Rhinovirus Subtype 16 PCR Kit (Primerdesign®) was used to quantify BEC viral load at 2, 4, 7, and 10 days post infection.

### RNA collection

To collect RNA from BECs in ALI, basolateral media was removed, and 800µL of lysis buffer (Invitrogen®, Product No. AM1912) was added to the apical surface of cultures. To mechanically disrupt BECs from the transwell membrane, the apical surface was gently scratched in a crosshatch pattern using a pipet tip. RNAqueous™ kit for total RNA isolation (Thermo Fisher Scientific) was used to extract RNA. This was performed prior to infection and at days 2, 4, 7, and 10 days post infection.

### Statistical Analysis

To identify genes with differential expression trajectories among the three treatment groups (RV16 Alone, IL-13 Alone, and IL-13 + RV16) across five timepoints (0, 2, 4, 7, and 10 days), a generalized additive mixed model (GAMM) was run with an interaction between time and treatment group and a random effect for epithelial cell donorID. This was done using the R package “mgcv” with model syntax of gene_expression ∼ treatment_group + s(time, k=5, bs=“cr”) + s(time, k=5, bs=“cr”, by=treatment_group) + s(donorId, bs=“re”). This model allowed the overall effect of time and the interaction between time and treatment group to be modeled flexibly using cubic regression splines, while accounting for repeated measures from the same donor through a random effect.

Predicted expression values for each gene were obtained using the “tidymv” package with the predict_gam() function, excluding the donor random effect to extract group-level smooth trajectories. The fitted values were scaled per gene, and hierarchical clustering was performed using Euclidean distance and complete linkage to identify groups of genes with similar modeled temporal trajectories. The clustering dendrogram was cut at a fixed height (h = 100), resulting in 26 gene clusters. These clusters represent groups of genes showing similar dynamic responses to IL-13 and/or viral infection over time.

Cluster expression values were summarized by taking the mean of all log2-transformed gene expression values belonging to each cluster for every sample. Using the same GAMM structure described above, these clusters were then modeled to identify those showing differential expression patterns over time among the three treatment groups at a false discovery rate (FDR) of 0.05 for either the group or interaction term. Multiple testing correction was performed using the Benjamini–Hochberg procedure. Healthy (n=11) and asthma (n=50) samples were combined for these analyses due to a lack of significant DEGs between groups with an FDR<0.05.

For clusters showing significant temporal differences among groups, pathway enrichment analysis was performed using the “clusterProfiler” package. Enrichment was tested using the gene ontology biological processes (GO_BP), KEGG, Reactome, Biocarta, and MSigDB Hallmark gene sets. A hypergeometric test was used to assess overrepresentation, and pvalues were adjusted for multiple testing using the Benjamini–Hochberg method. Pathways were considered significantly enriched at an FDR 0.05. The most significantly enriched pathways were used to annotate clusters and aid biological interpretation.

To assess the association between cluster expression and viral load, linear mixed effects models were run using the “kimma” package with model syntax: cluster_expression ∼ log10(viral load) + time + (1|donorId). Models were fit across post-infection samples, and significant associations between cluster expression and viral load were identified at an FDR 0.05.

## RESULTS

Primary BECs were collected from pediatric patients with (n=50) and without (n=11) asthma (Table 1). There were no significant differences in age, sex, or race/ethnicity between asthma and healthy donors. Total serum IgE was higher in donors with asthma compared with healthy controls (p<0.01). Donors with asthma demonstrated sensitivity to aeroallergens as measured by antigen-specific IgE, whereas healthy donors did not demonstrate any sensitivity to tested aeroallergens. FEV1/FVC was lower in the asthma donors as compared to healthy donors (p=0.01). 54% of donors with asthma were on inhaled corticosteroids at the time of BEC collection. No donors were on biologic therapies for asthma.

**TABLE 1.** BEC donor characteristics.

|  | **Asthma Donors**<br>**(n=50)** | **Healthy Donors**<br>**(n=11)** |
| --- | --- | --- |
| Age<br>(yrs., mean ± SD) | 11.7 ± 4.0 | 12.9 ± 3.6 |
| Female Sex | 22 (44%) | 5 (45%) |
| American Indian or<br>Alaska Native | 1 (2%) | 0 (0%) |
| Asian | 2 (4%) | 1 (9%) |
| Black | 4 (8%) | 1 (9%) |
| Native Hawaiian or<br>Pacific Islander | 1 (2%) | 0 (0%) |
| White | 38 (76%) | 9 (82%) |
| Hispanic | 6 (12%) | 0 (0%) |
| Not Hispanic | 44 (88%) | 11 (100%) |
| Serum IgE<br>(mean ± SD) | 254.4 ± 401.4 | 24.8 ± 30.2 |
| $\geq 2$ aeroallergen<br>positive by allergen-specific IgE | 25 (50%) | 0 (0%) |
| FeNO<br>(ppb; mean ± SD) | 15.8 ± 16.4 | 11.4 ± 8 |
| FEV<sub>1</sub> % predicted<br>(mean ± SD) | 100.1 ± 14 | 103.5 ± 17 |
| FEV<sub>1</sub>/FVC<br>(mean ± SD) | 0.83 ± 0.06 | 0.87 ± 0.04 |
| Inhaled corticosteroid<br>(ICS) use at time of<br>BEC collection | 27 (54%) | N/A |
| History of Severe<br>Asthma Exacerbation | 30 (60%) | N/A |
| Body Mass Index<br>(mean ± SD) | 23.8 ± 8.1 | 20.8 ± 5.2 |

BEC = Bronchial epithelial cell
SD = Standard deviation
IgE = Immunoglobulin
FeNO = Fractional exhaled nitric oxide
FEV<sub>1</sub> = Forced expiratory volume in 1 second
FVC = Forced vital capacity
BMI = Body mass index

### IL-13 pretreatment alters antiviral effector programs and secondary pro-inflammatory responses while preserving upstream interferon induction in virus-infected bronchial epithelial cells

Bronchial epithelial cells (BECs) were differentiated at an air-liquid interface for 21 days into a confluent pseudostratified epithelium (Figure 1A-B). To model type-2 inflammation, differentiated BECs were treated with IL-13 (10ng/mL) for 7 days prior to infection with RV-A16 (MOI 0.5) (Figure 1A). We observed histologic changes in the BECs in ALI following IL-13 treatment. Specifically, expected goblet cell hyperplasia was observed via hematoxylin and eosin (H&E) and immunofluorescent MUC5AC staining (Figure 1B). There was also a visible increase in patches within the epithelium via scanning electron microscopy (SEM) (Figure 1C) that represented areas of goblet cell degranulation of mucus observed via transmission electron microscopy (TEM) (Figure 1D).

**Figure 1.**
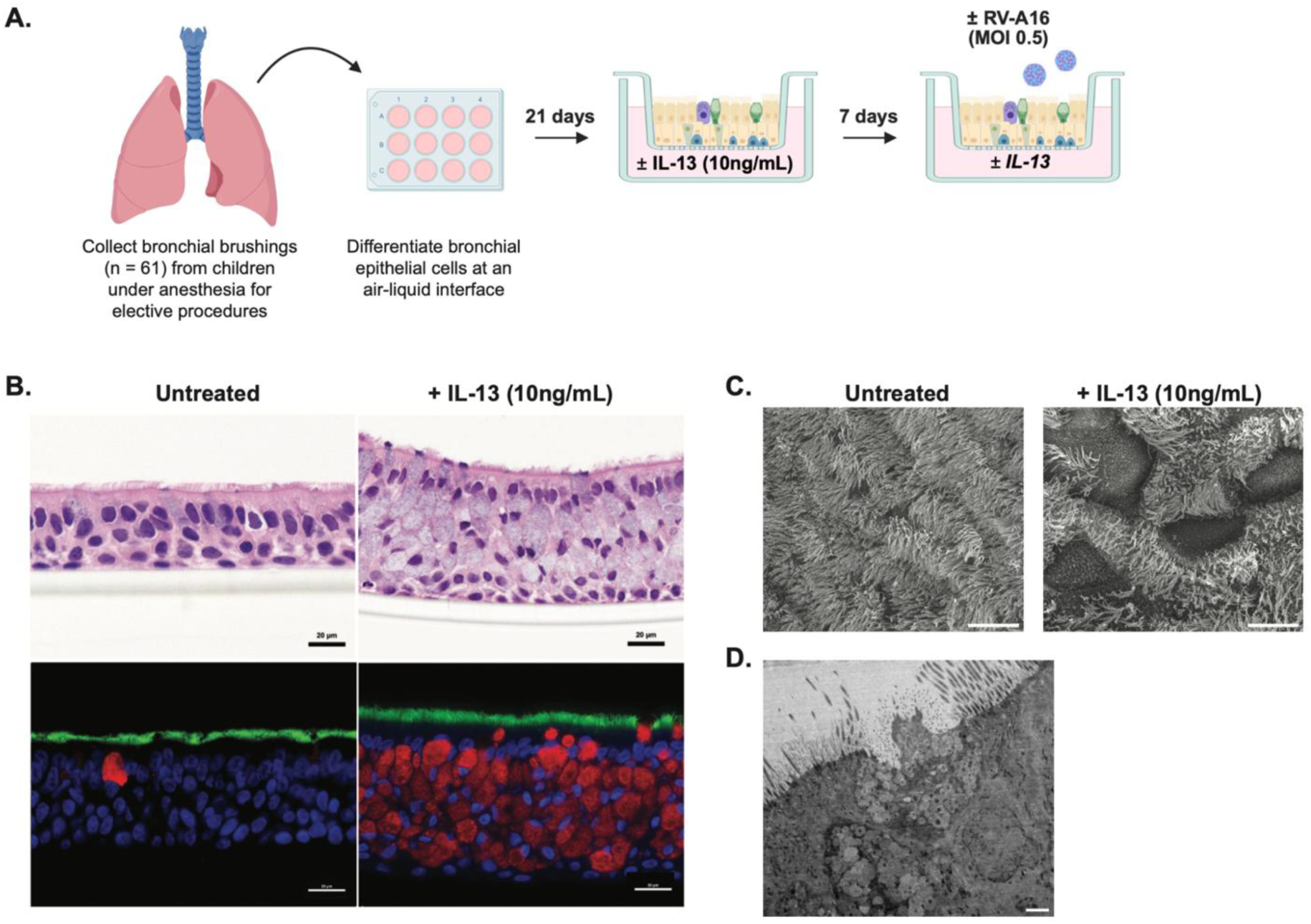
Experimental design and modeling of T2 inflammation. (A) Schematic of the experimental design. Bronchial brushings were collected from children (n=61) under anesthesia for elective procedures. Cells were differentiated at an air-liquid interface culture for 21 days, then treated with IL-13 (10ng/mL) for 7 days prior to apical infection with human rhinovirus A16 (RV-A16) at an MOI 0.5. IL-13 treatment was continued through the infection timepoints. (B) H&E (top panels) and immunofluorescence (bottom panels) microscopy of bronchial epithelial cells in air-liquid interface cultures untreated and treated with IL-13 (10ng/mL). Immunofluorescence key: Blue = DAPI, Green = Tubulin, Red = MUC5AC. (C) Scanning electron microscopy (SEM) images of BECs untreated and treated with IL-13 (10ng/mL). (D) transmission electron microscopy (TEM) image of goblet cell degranulating mucus.

Epithelial gene expression was measured prior to infection and at days 2, 4, 7, and 10 post infection. In response to IL-13 stimulation in the absence of rhinovirus, BECs displayed 10253 DEGs (Figure 2A), including leading edge canonical response genes *POSTN* and *SERPINB2*, as well as *MUC5AC* indicative of goblet cell hyperplasia (Supplemental Figure 1). In response to rhinovirus infection alone, BECs displayed 7727 differentially expressed genes (DEGs) (FDR <0.05) with leading edge genes related to interferon responses (Figure 2B). Notably, 7095 genes showed an interaction effect between RV infection and IL-13 stimulation (Supplemental Table 1).

**Figure 2.**
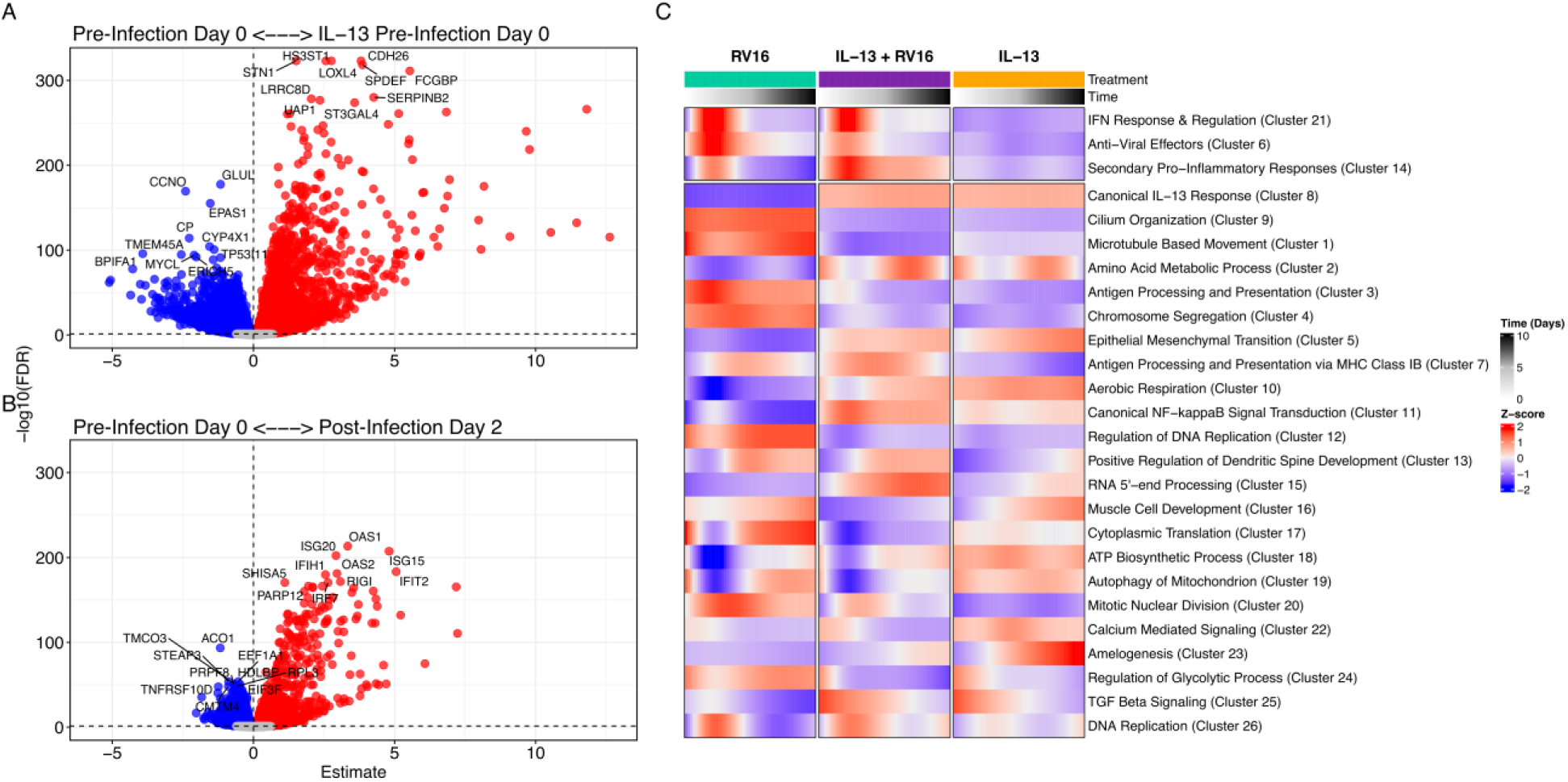
IL-13 stimulation and RV16 infection drive distinct transcriptional programs, and GAM-based clustering reveals coordinated temporal response patterns. (A) Volcano plot showing differential gene expression between IL-13–treated vs untreated epithelial cultures at pre-infection Day 0. Red points indicate significantly upregulated genes (4838 genes) and blue points indicate significantly downregulated genes (5415 genes). (B) Volcano plot showing differential gene expression between post-Infection Day 2 vs untreated Day 0 epithelial cultures at pre-infection. Red points indicate significantly upregulated genes (3482 genes) and blue points indicate significantly downregulated genes (4245 genes). FDR threshold of 0.05 shown by dashed line. Estimates represent log fold-changes from the linear model. C) The heat map shows the mean GAM predicted values of the 26 Clusters. Mean GAM predicted values levels are shown as row normalized Z-scores for each group and timepoint (columns) with red representing higher relative expression and blue representing lower relative expression. The 3 clusters of interest are presented at the top. Those clusters are named according to their annotation summary. The other clusters are labeled by their top pathway enrichment score. The number in parentheses indicates the cluster number.

Hierarchical clustering of GAMM fit models revealed clusters that had shared temporal expression patterns (Figure 2C). Focusing on our hypothesis that IL-13 negatively affects antiviral interferon responses, we specifically investigated three separate gene clusters enriched in interferon response genes (ISGs) (Figure 3A, Supplemental Table 2). The first such cluster contained type-I and type-III interferon genes and upstream regulators of the interferon response, annotated as interferon antiviral response and regulation (Figure 3B, Supplemental Table 3). It consisted of 313 total genes including 15 cytokines and 40 transcription factors (IL13 + RV16 vs RV16 GAMM Shape: FDR=3.4e-04). Notable genes included *IFNB1, IFNL1, IFNL2, IFNL3, CXCL10, CXCL11, IRF7, ETV7, STAT1, STAT2, GBP5, TLR2, and TLR3*. Expression of this cluster peaked at 2d post infection and showed statistically similar expression patterns with or without IL-13 stimulation, suggesting that IL-13 does not affect expression of interferons and the upstream elements of the ISG cascade.

**Figure 3.**
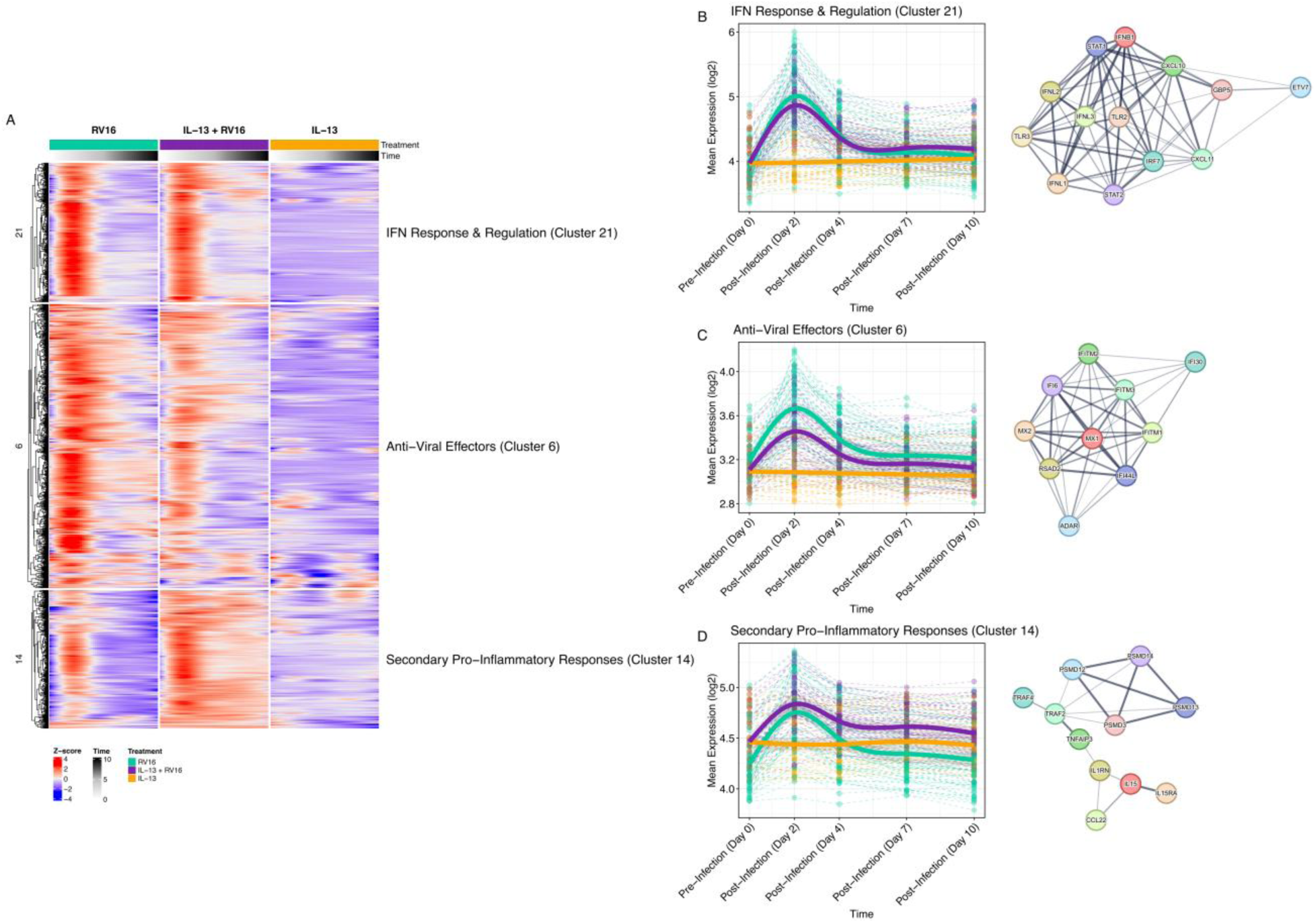
IL-13 pretreatment alters antiviral effector programs and secondary proinflammatory responses while preserving upstream interferon induction in virus-infected bronchial epithelial cells. A) Heatmap showing the temporal expression patterns of three interferon-associated gene clusters identified by hierarchical clustering of GAMM-fitted expression trajectories. Rows represent genes, and columns represent timepoints (0, 2, 4, 7, and 10 days) across the three treatment conditions (RV16, IL-13 + RV16, IL-13). Row values are displayed as Z-scores of mean predicted expression, with red indicating higher relative expression and blue indicating lower expression. Clusters displayed include IFN Response & Regulation (Cluster 21), Anti-Viral Effectors (Cluster 6), and Secondary Pro-Inflammatory Responses (Cluster 14). Annotations on the right summarize the dominant biological processes enriched within each cluster. B) GAMM plot showing the non-linear changes in expression of the IFN Response & Regulation cluster (Cluster 21) across treatments and timepoints. Expression peaked at Day 2 post-infection and showed comparable temporal trajectories across RV16 and IL-13 + RV16 groups (IL-13 + RV16 vs RV16: GAMM Shape: FDR=3.4e-04). C) GAMM plot showing the expression dynamics of the Anti-Viral Effectors cluster (Cluster 6) across treatments and timepoints. Expression peaks at Day 2 post-infection with the RV16 group showing a higher expression at all post-infection timepoints (IL-13 + RV16 vs RV16: GAMM Shape: FDR=1.8e-04). D) GAMM plot illustrating temporal expression of the Secondary ProInflammatory Responses cluster (Cluster 14). Expression peaked at Day 2 post-infection with a higher expression in IL-13 + RV16 group, with the largest differences emerging at later postinfection timepoints (IL-13 + RV16 vs RV16: GAMM Shape: FDR=<1.0e-10). String networks are showing representative genes for the clusters.

In contrast, the second cluster showed greater enrichment of anti-viral effector molecules, which were significantly decreased by IL-13 pretreatment in rhinovirus infection (IL-13 + RV16 vs RV16 GAMM Shape: FDR=1.8e-04) (Figure 3C, Supplemental Table 3). This cluster peaked at 2 days post infection but showed a lower magnitude of expression across all post infection timepoints due to IL-13 pretreatment. It was composed of 638 genes including effectors molecules that limit viral replication and spread such as *MX1, MX2, RSAD2, IFITM1, IFITM2, IFITM3*, as well as multiple additional IFI family genes. Conversely, IL-13 preconditioning in rhinovirus infection increased the expression of a gene cluster that we annotated as “secondary proinflammatory response” (IL-13 + RV16 vs RV16 GAMM Shape: FDR=<1.0e-10) (Figure 3D, Supplemental Table 3). This gene cluster was composed of 311 genes related to a mixed immune response including the IL-15 pathway, complement, antigen presentation, endoplasmic reticulum stress, cell death pathways, and TNF signaling. Genes included *IL15, IL15RA, C2, IL1RN, CCL22, PSMD* family members, *TAP1/2, TNFAIP3, and TRAF2/TRAF4.* Gene expression differences between BECs that were and were not preconditioned with IL-13 were most prominent at later time points after viral infection.

### IL-13 preconditioning of the bronchial epithelium did not significantly alter viral load

We next examined the effect of IL-13 preconditioning on viral load. Rhinovirus viral load was measured by PCR at days 2, 4, 7, and 10 post infection. When comparing BECs with and without IL-13 preconditioning, there were no significant differences in viral load across any timepoint (Figure 4). This was in the setting of heterogeneous responses to IL-13 among different donors. Similar viral load across conditions (p=1.0) suggested the gene expression differences observed with and without IL-13 preconditioning stemmed from an IL-13 and RV interaction rather than a group level difference in viral load.

**Figure 4.**
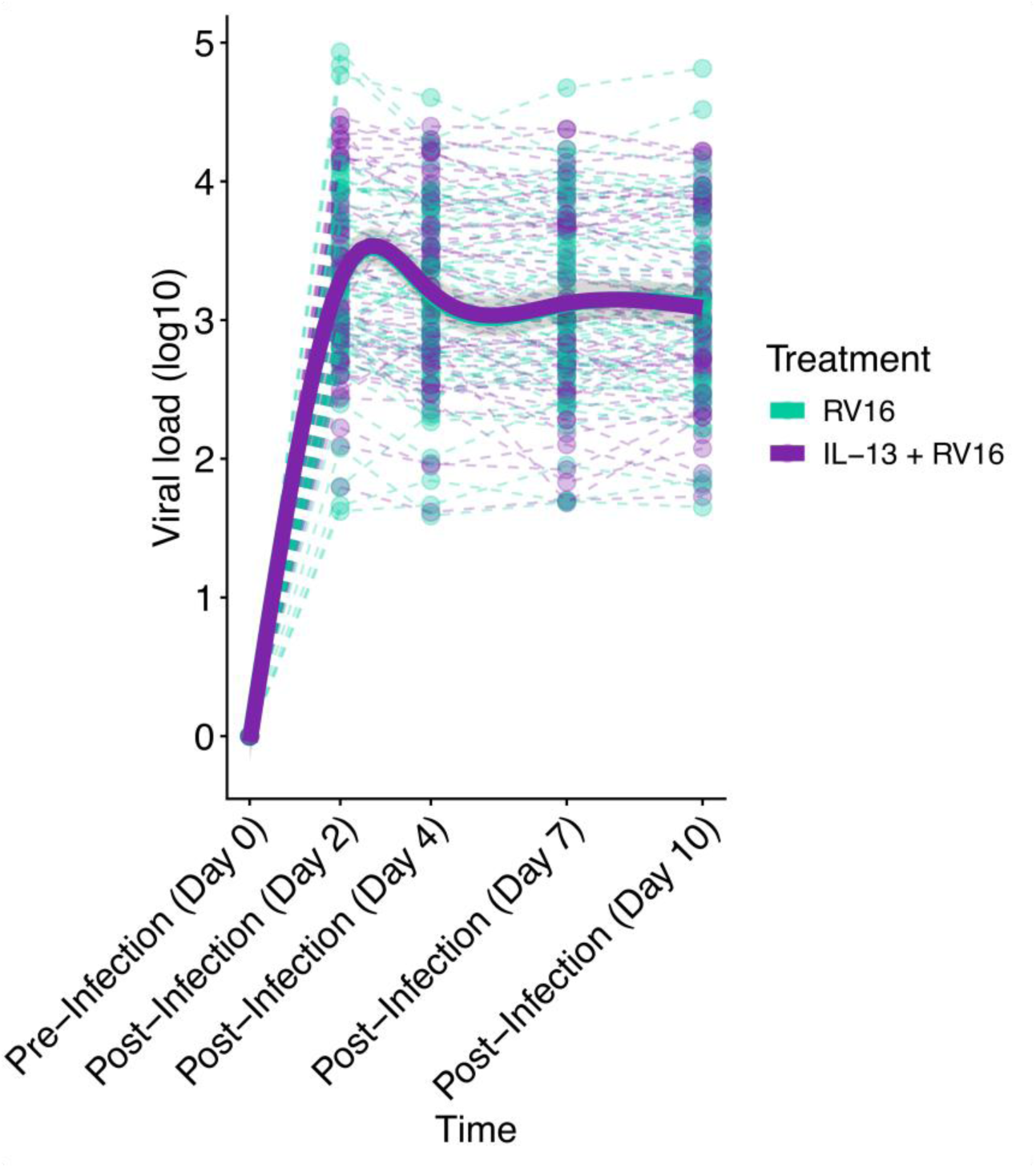
IL-13 preconditioning of the bronchial epithelium did not alter viral load across timepoints. GAMM plot showing the non-linear patterns in viral load over RV16 and IL-13 + RV16 groups (IL-13 + RV16 vs RV16: GAMM Shape: p=1.0). Fit lines are based on a generalized additive mixed model including 95% confidence intervals.

### IL-13 decouples viral load from epithelial immune responses in RV infection

When we examined the association of viral load on the IFN Response & Regulation, Anti-Viral Effector, and Secondary Proinflammatory Response gene clusters, we found strong positive correlations between viral copy number and mean cluster expression (Estimates 0.15-0.31, p< 1e-04, Figure 5A,C,E, Supplemental Table 4). IL-13 preconditioning of the epithelium eliminated these associations (Figure 5B,D,F), suggesting that while IL-13 preconditioning did not alter viral load, it may dysregulate expected viral immune responses.

**Figure 5.**
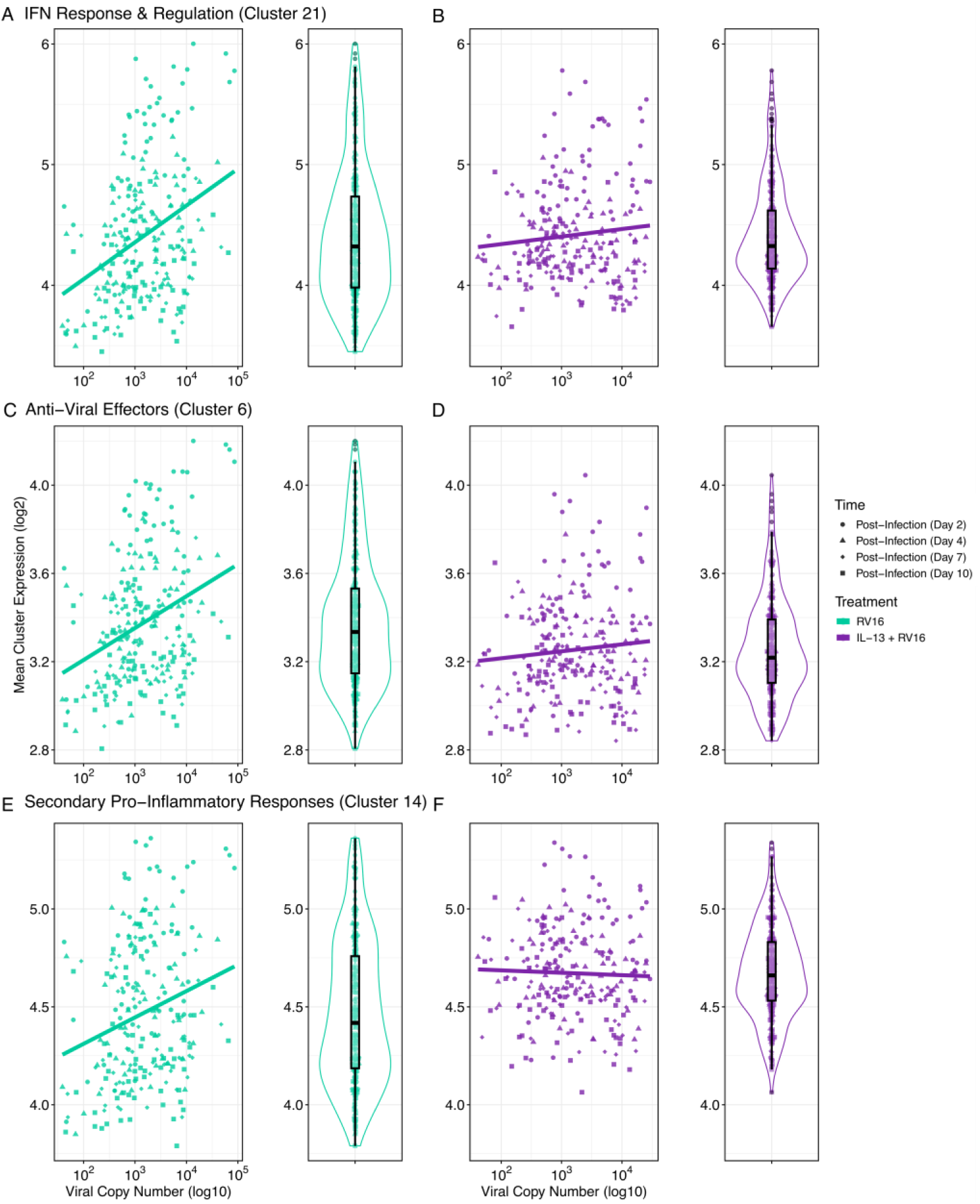
IL-13 preconditioning decouples viral load from epithelial immune responses in RV infection. (A–F) Scatterplots and corresponding distribution plots showing the association between viral copy number (log10) and mean expression (log2) of three key epithelial gene clusters identified from the GAMM analysis: IFN Response & Regulation (Cluster 21), Anti-Viral Effectors (Cluster 6), and Secondary Pro-Inflammatory Responses (Cluster 14). A, C, and E show relationships in untreated RV16-infected BECs, whereas B, D, and F show the same relationships in IL-13 preconditioned + RV16-infected BECs.

## DISCUSSION

The impact of IL-13 on rhinovirus infection and innate immune and inflammatory responses to rhinovirus infection by the bronchial epithelium in asthma is poorly understood. There are mixed reports describing increased (32) and decreased (33) susceptibility to rhinovirus infection due to IL-13-induced mucous cell hyperplasia. Here, we leverage primary BECs from children with and without asthma that were differentiated to an organotypic state at an air liquid interface to show that type 2 inflammation, as modeled by high IL-13 conditioning (23,24), may not affect viral load, but rather alters other subsequent anti-viral and inflammatory processes that are relevant to the pathogenesis of viral-triggered asthma exacerbation.

Studies by Jayavelu and Benson et al. and by Gaberino et al. described that low baseline interferon expression in the bronchial and nasal epithelium, respectively, was associated with more pronounced increases in interferon responses during respiratory illness, increased viral load, and increased exacerbation risk in pediatric asthma (18,20). Paradoxically, type 2 inflammation has been shown to decrease SARS-CoV2 replication in the asthmatic epithelium (18,34) through downregulation of functional ciliated epithelial pathways, specifically those governing differentiation, as well as the production and function of cilia and axonemes. In this study we hypothesized that T2 inflammation, a common driver of pediatric asthma, would decrease the expression of interferon responses to rhinovirus infection, increase viral load, and increase post-infection inflammatory pathways. Demonstrating a degree of internal validity, our data corroborate existing literature that describe bronchial epithelial IFN responses to respiratory viral infection (16,20,34,35,36), activation of canonical T2 genes such as *SERPINB2* and *POSTN* by IL-13 (37), and IL-13 induction of *MUC5AC* (37,38) with accompanying histologic changes. However, pretreatment of BECs with IL-13 prior to rhinovirus infection did not alter viral load and did not affect a gene cluster associated with interferon response and regulation. This included viral sensors like TLRs, interferon regulatory factors such as STAT1/2, interferons (IFNB1, IFNL1, IFNL2, IFNL3), and ISGs like CXCL10 and CXCL11. Lack of changes in viral load and this gene cluster may be a product of donor heterogeneity, but taken together, data suggest that high T2 inflammation does not dampen these interferon response and regulation genes. We did, however, identify two other gene clusters that provide novel insight into epithelial responses driven by the interaction of IL-13 and rhinovirus.

One of these clusters was decreased by IL-13 and enriched for anti-viral effectors separate from the interferon response and regulation cluster. It included genes such as *MX1, MX2, RSAD2, IFITM1, IFITM2, IFITM3*, and multiple IFI family proteins. There is certainly overlap with some interferon-induced anti-viral responses, however, this finding argues that type 2 inflammation may decrease distinct anti-viral responses. While IL-13 did not alter viral load, it may impact subsequent immune responses and clinical sequelae. For example, this cluster included *IFITM1* and *IFITM3.* In a clinical study by Ravi et al, expression of IFITM1 and IFITM3 (but not other interferon response genes) in the bronchial epithelium of asthmatic patients was inversely correlated with subject decline in FEV1 after rhinovirus challeng*e* (39).

IL-13 increased expression of a cluster we annotated as “secondary proinflammatory responses”. This included genes associated with the IL-15, endoplasmic reticulum stress, cell death, and TNF signaling pathways. IL-15 is involved in T and NK cell homeostasis (40,41). In a mouse model of asthma, viral PAMP-induced exacerbations were dependent on presence of intact IL-15 (40). A recent study by Ramonell et al (42) showed that airway IL-15 expression was increased in patients with severe asthma as compared to patients with mild to moderate asthma. *IL15* expression also correlated with decreased lung function as measured by FEV1 (42). TNF signaling has been implicated in airway hyperresponsiveness and airway inflammation in asthma (43–45). Changes in genes associated with ER stress and cell death pathways suggest an effect of type 2 inflammation on responses to epithelial damage in asthma, which could affect remodeling that is known to be abnormal in patients with asthma (6,31). These gene expression differences between IL-13 treated and untreated conditions were most prominent at later time points in the infection course, suggesting that high type 2 inflammation prior to rhinovirus infection may promote more inflammatory late responses to rhinovirus infection that could have clinical consequences in pediatric patients with type 2 asthma.

We showed that these three clusters—Interferon Response & Regulation, Anti-Viral Effectors, and Secondary Proinflammatory Responses, all correlated with viral load in the absence of IL13 pretreatment. Elimination of that correlation by pre-treatment with IL-13 suggests that high T2 inflammation disrupts typical epithelial immune responses.

Our data are limited by a modest healthy control sample size (n=11). We suspect that the lack of observed differences in gene expression between asthma and healthy groups may be due to both donor heterogeneity and sample size such that we are not adequately powered to detect differences between healthy and asthma groups. Additionally, it is known that asthma consists of multiple endotypes (5,46), and T2 asthma is one of those endotypes common in pediatric asthma. In this study, we do not delineate between molecular endotypes. IL-13 pretreatment of all donor samples modeled a T2 environment but may not account for already existing differences in the donor epithelium due to their asthma endotype or other potential differences including epigenetic changes.

Future studies will include recruitment of additional healthy donors to assess differential responses between healthy and asthma samples. Our group previously reported that IL-13 treatment of bronchial epithelial cells reduced SARS-CoV2 replication in donors with allergic asthma and not healthy children (34), suggesting there may be differential responses to IL-13 in rhinovirus between healthy and asthmatic epithelium. Although non-type 2 asthma is less common among children, in future studies we plan to investigate differences in RV replication and BEC responses to viral infection between pediatric asthma donors with type 2 vs. non-type 2 asthma.

Our findings refine the mechanistic framework through which anti–IL-4Rα biologics, such as dupilumab, may influence viral susceptibility and post-viral inflammation in T2-high asthma. IL4Rα–targeting antibodies broadly suppress IL-13-driven type 2 epithelial programs, including mucus-associated gene expression and airway remodeling signatures (47), and experimental IL-4Rα blockade in bronchial epithelial cells reduces T2 cytokine–driven mediators such as CCL26 and TSLP while leaving antiviral IFNβ and IFNλ1 responses largely intact during rhinovirus infection (48). Our data showing that IL-13 selectively attenuates epithelial antiviral effector clusters (e.g., MX1/2, RSAD2, IFITM1/2/3) while augmenting secondary inflammatory pathways suggest that IL-4Rα blockade may not primarily enhance canonical interferon induction but rather may restore the balance between epithelial antiviral competence and downstream inflammatory amplification. This distinction may be clinically meaningful, as the antiviral effector pathways suppressed by IL-13 in our study include genes previously linked to airflow obstruction and epithelial injury during viral infections (e.g., IFITM1/3)(39). Therefore, our data support a model in which dupilumab’s clinical benefit in reducing exacerbations may arise not only by moderating chronic T2 inflammation, but also by preventing IL-13 from skewing epithelial responses toward a dysregulated, proinflammatory state upon viral exposure.

These observations provide novel mechanistic insights into why some T2-high patients remain susceptible to viral-triggered exacerbations even with IL-4Rα blockade, identifying epithelial pathways that may underlie persistent risk. Although IL-13 induced large-scale changes in epithelial gene expression, it did not suppress interferon-regulatory genes or alter viral load, suggesting that defects in upstream interferon induction may persist independently of IL-13 signaling. This aligns with recent evidence that low baseline/pre-infection epithelial interferon tone itself is an asthma endotype associated with high viral replication and exacerbation risk (18,20,39). Second, the IL-13-induced secondary inflammatory cluster–enriched for IL-15, TNF-NFκB signaling, ER stress, and cell-death pathways–was most strongly induced at later infection timepoints. These pathways have been independently linked to severe exacerbations (42,45) and may remain operative even when IL-13 signaling is therapeutically suppressed. Together, these data suggest a model in which persistent epithelial interferon insufficiency combined with non-IL-13 inflammatory programs (TNF/ER-stress/cell-death) may drive breakthrough viral exacerbations in biologic-treated T2-high patients. Targeting these mechanisms, through agents that enhance epithelial antiviral effector function, modulate ERstress responses, or inhibit TNF-family signaling, may represent promising adjunctive therapeutic strategies.

In summary, this study demonstrates that IL-13 profoundly reshapes bronchial epithelial responses to rhinovirus not by altering viral load or interferon induction, but by suppressing antiviral effector programs and amplifying proinflammatory injury pathways. These findings provide a mechanistic basis linking T2 inflammation to severe viral-triggered exacerbations in pediatric asthma, while also revealing why IL-13–targeting therapies may be effective for many but not all patients. By identifying epithelial pathways that remain vulnerable even in the presence of IL-13 blockade, this study identifies new potential therapeutic targets for children with T2-high asthma who continue to experience viral-triggered exacerbations despite biologic therapy.

## ACKNOWLEDGEMENTS

We would like to acknowledge the BRI Genomics Core, RRID:SCR_026658 for use of their facilities for generating and processing next-generation sequencing data.

## Abbreviations used

ALI: Air liquid interface
BEC: Bronchial epithelial cell
IgE: Immunoglobulin E
FeNO: Fraction of exhaled nitric oxide
FEV_1_: Forced expiratory volume in 1 second
FVC: Forced vital capacity
RV: Human Rhinovirus
IFN: Interferon
SD: Standard deviation

## SUPPLEMENTAL INFORMATION

### Supplemental Methods

#### RNA Sequencing and Data Processing

The SMART-Seq v4 Ultra Low Input RNA Kit for Sequencing (Takara, San Jose, Calif) was used to create libraries from total RNA. Libraries were reverse transcribed then amplified to generate full-length cDNA amplicons. The NexteraXT DNA sample preparation kit with unique dual indexes (Illumina, San Diego, Calif) created Illumina-compatible barcoded sequencing libraries. A Qubit Fluorometer (Thermo Fisher Scientific, Waltham, Mass) pooled and quantified libraries with subsequent sequencing of pooled libraries on a NextSeq 2000 sequencer (Illumina). Sequencing was done with paired-end 53-base reads and a target depth of 5 million reads per sample. BaseSpace (Illumina) was used to process base calls and quality-trim lowconfidence base calls from read ends. Casava (Illumina) was used to deconvolute and convert resulting bcl files to fastq files that were subsequently aligned to the Ensembl human genome (GRCh38, Ensembl 91) via STAR (version 2.4.2a). Gene counts were generated with HTSeqcount (version 0.4.1) using the parameters of “intersection (nonempty)” mode, minimum alignment quality of 20, and otherwise default parameters. PICARD (version 1.134), FASTQC (version 0.11.3), Samtools (version 1.2), and HTSeq-count (version 0.4.1) were used to generate quality metrics. Samples were used if quality metrics showed human aligned counts greater than 1 million mapped reads and a median coefficient of variation coverage less than 0.7. A gene filter was applied to include only protein coding genes that had a trimmed mean of M value normalization count of at least 0.3 in at least 10% of samples. The limma R package (version 4.2.1) function voomWithQualityWeights was used to transform normalized counts to log2 counts per million mapped reads along with observations level weights.

#### Data Availability

GSE for RNA sequencing data upload to NCBI Gene Expression Omnibus database is pending.

#### Histopathology

Hematoxylin and eosin (H&E) staining and immunofluorescence were performed on 10% formalin-fixed paraffin-embedded 5-µm sections from transwell membranes oriented en face. Dual immunofluorescence for rabbit alpha tubulin (1:1000 dilution; Invitrogen, #PA5-105102) and mouse MUC5AC (1:300 dilution; ab3649, Abcam) was carried out following citrate pH 6.0 antigen retrieval and serum block; antibodies were incubated for 2 hours at room temperature. Alpha tubulin was developed with donkey anti-rabbit Alexa Fluor 488 and MUC5AC with donkey antimouse cyanine (Cy3) (both 1:1000 Jackson ImmunoResearch). Coverslips were mounted using Vectashield fluorescent mounting medium with DAPI (Vector Laboratories). Images were visualized and captured with a digital camera mounted on a Nikon Eclipse 80i microscope using NIS-Elements Advanced Research Software 6.10.01 (Nikon Instruments Inc., Melville, NY). For scanning electron microscopy (SEM), BECs in transwells were fixed in 2% paraformaldehyde and 2% glutaraldehyde in 0.1M phosphate buffer at 4C°. 0.1M phosphate was used for rinsing and samples were post-fixed in 1% osmium tetroxide overnight at 4C°. Samples were then washed in water, dehydrated with 100% ethanol, and critical point dried. Gold sputter coating was then applied. Samples were imaged using a Thermo Fisher Scientific FEI Apreo VolumeScope scanning electron microscope.

For transmission electron microscopy (TEM), transwells were fixed and dehydrated in the same manner as samples processed for SEM. Briefly, transwells were fixed in 2% paraformaldehyde and 2% glutaraldehyde in 0.1M phosphate buffer and post-fixed in 1% osmium tetroxide, then gradually dehydrated to 100% EtOH. After dehydration, transwell membranes were removed with a biopsy punch tool and transitioned through propylene oxide, gradually infiltrated with Spurr’s low viscosity resin, until finally embedding and curing at 70C° for 24 h. Resin blocks were sectioned with Leica EM UC7 Ultramicrotome at 70nm using a glass knife on formvarcoated 100-mesh copper grids. Samples were then stained with 2% uranyl acetate for 8 min, Reynolds lead citrate for 5 min, then imaged using the FEI Tecnai G2 20 TEM.

### Supplemental Figures

**Supplemental Figure 1.**
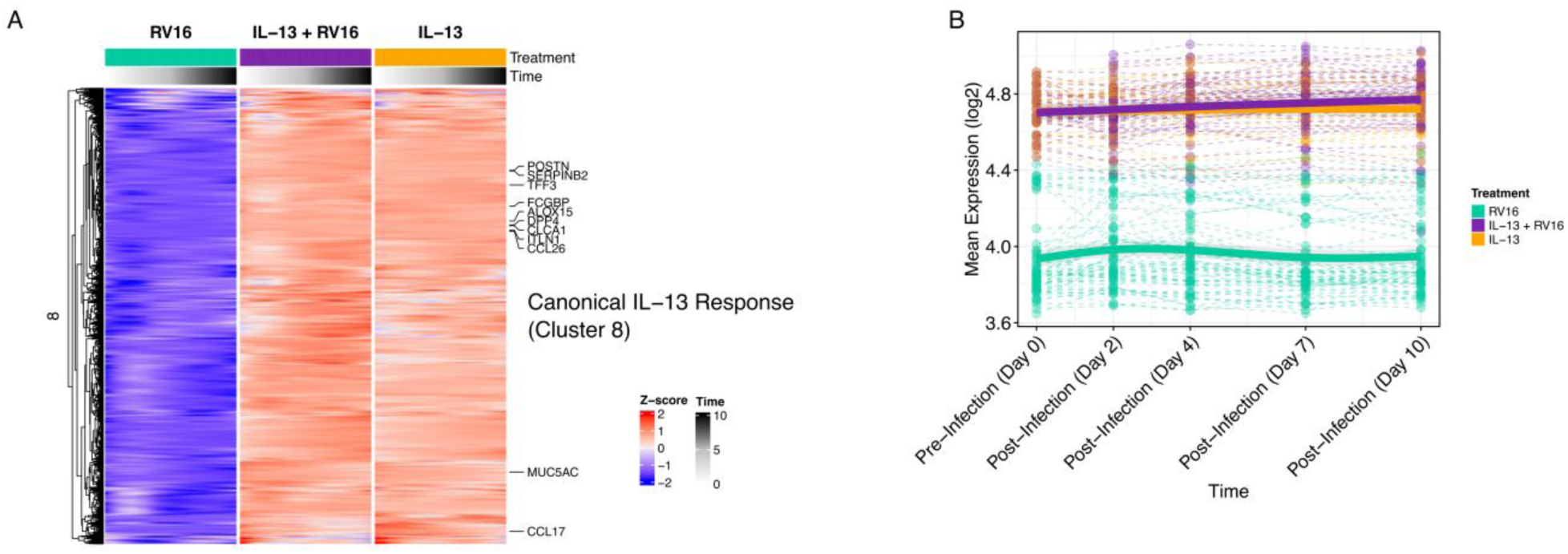
Induction of IL-13 canonical response genes. A) Heatmap of the cluster annotated as Canonical IL-13 Response identified by hierarchical clustering of GAMMfitted expression trajectories. Rows represent genes, and columns represent timepoints (0, 2, 4, 7, and 10 days) across the three treatment conditions (RV16, IL-13 + RV16, IL-13). Row values are displayed as Z-scores of mean predicted expression, with red indicating higher relative expression and blue indicating lower expression. Notable IL-13-induced genes are labeled. B) GAMM plot showing the non-linear changes in expression of the Canonical IL-13 Response cluster (cluster 8) across treatments and timepoints.

### Supplemental Tables

**Supplemental Table 1.** Results of the Linear Model showing the differential expressed genes by the treatment groups

**Supplemental Table 2.** Table of genes in Clusters with Cluster annotations

**Supplemental Table 3:** Generalized Additive Mixed Models (GAMMs) results comparing expression differences over time between IL-13 treatment groups. Table shows the smoothing terms of time and smoothing interaction term for the effect of IL-13 groups over time.

**Supplemental Table 4:** Linear Model comparison of viral copy number to the cluster expression.

## REFERENCES

1. Nurmagambetov T, Kuwahara R, Garbe P. The Economic Burden of Asthma in the United States, 2008-2013. Ann Am Thorac Soc. 2018 Mar;15(3):348–56.

2. Jartti T, Gern JE. Role of viral infections in the development and exacerbation of asthma in children. J Allergy Clin Immunol. 2017 Oct;140(4):895–906.

3. Khetsuriani N, Kazerouni NN, Erdman DD, Lu X, Redd SC, Anderson LJ, et al. Prevalence of viral respiratory tract infections in children with asthma. J Allergy Clin Immunol. 2007 Feb;119(2):314–21.

4. Maspero J, Adir Y, Al-Ahmad M, Celis-Preciado CA, Colodenco FD, Giavina-Bianchi P, et al. Type 2 inflammation in asthma and other airway diseases. ERJ Open Res [Internet]. 2022 Jul;8(3). Available from: 10.1183/23120541.00576-2021

5. Ray A, Das J, Wenzel SE. Determining asthma endotypes and outcomes: Complementing existing clinical practice with modern machine learning. Cell Rep Med. 2022 Dec 20;3(12):100857.

6. Lopez-Guisa JM, Powers C, File D, Cochrane E, Jimenez N, Debley JS. Airway epithelial cells from asthmatic children differentially express proremodeling factors. J Allergy Clin Immunol. 2012 Apr;129(4):990–7.e6.

7. Fahy JV. Type 2 inflammation in asthma--present in most, absent in many. Nat Rev Immunol. 2015 Jan;15(1):57–65.

8. Altman MC, Gill MA, Whalen E, Babineau DC, Shao B, Liu AH, et al. Transcriptome networks identify mechanisms of viral and nonviral asthma exacerbations in children. Nat Immunol. 2019 May;20(5):637–51.

9. Contoli M, Ito K, Padovani A, Poletti D, Marku B, Edwards MR, et al. Th2 cytokines impair innate immune responses to rhinovirus in respiratory epithelial cells. Allergy. 2015 Aug;70(8):910–20.

10. Bianco A, Whiteman SC, Sethi SK, Allen JT, Knight RA, Spiteri MA. Expression of intercellular adhesion molecule-1 (ICAM-1) in nasal epithelial cells of atopic subjects: a mechanism for increased rhinovirus infection? Clin Exp Immunol. 2000 Aug;121(2):339–45.

11. Cakebread JA, Haitchi HM, Xu Y, Holgate ST, Roberts G, Davies DE. Rhinovirus-16 induced release of IP-10 and IL-8 is augmented by Th2 cytokines in a pediatric bronchial epithelial cell model. PLoS One. 2014 Apr 4;9(4):e94010.

12. Jakiela B, Rebane A, Soja J, Bazan-Socha S, Laanesoo A, Plutecka H, et al. Remodeling of bronchial epithelium caused by asthmatic inflammation affects its response to rhinovirus infection. Sci Rep. 2021 Jun 17;11(1):12821.

13. Contoli M, Message SD, Laza-Stanca V, Edwards MR, Wark PAB, Bartlett NW, et al. Role of deficient type III interferon-lambda production in asthma exacerbations. Nat Med. 2006 Sep;12(9):1023–6.

14. Wark PAB, Johnston SL, Bucchieri F, Powell R, Puddicombe S, Laza-Stanca V, et al. Asthmatic bronchial epithelial cells have a deficient innate immune response to infection with rhinovirus. J Exp Med. 2005 Mar 21;201(6):937–47.

15. Spann KM, Baturcam E, Schagen J, Jones C, Straub CP, Preston FM, et al. Viral and host factors determine innate immune responses in airway epithelial cells from children with wheeze and atopy. Thorax. 2014 Oct;69(10):918–25.

16. Altman MC, Reeves SR, Parker AR, Whalen E, Misura KM, Barrow KA, et al. Interferon response to respiratory syncytial virus by bronchial epithelium from children with asthma is inversely correlated with pulmonary function. J Allergy Clin Immunol. 2018 Aug;142(2):451– 9.

17. Bhakta NR, Christenson SA, Nerella S, Solberg OD, Nguyen CP, Choy DF, et al. IFN-stimulated Gene Expression, Type 2 Inflammation, and Endoplasmic Reticulum Stress in Asthma. Am J Respir Crit Care Med. 2018 Feb 1;197(3):313–24.

18. Gaberino CL, Altman MC, Gill MA, Bacharier LB, Gruchalla RS, O’Connor GT, et al. Dysregulation of airway and systemic interferon responses promotes asthma exacerbations in urban children. J Allergy Clin Immunol. 2025 May;155(5):1499–509.

19. Patel DA, You Y, Huang G, Byers DE, Kim HJ, Agapov E, et al. Interferon response and respiratory virus control are preserved in bronchial epithelial cells in asthma. J Allergy Clin Immunol. 2014 Dec;134(6):1402–12.e7.

20. Doni Jayavelu N, Benson B, Dela Cruz PC, Powell WT, Rich LM, Vanderwall ER, et al. Bronchial Epithelial Transcriptome Reveals Dysregulated Interferon and InflammatoryResponses to Rhinovirus in Exacerbation-Prone Pediatric Asthma. JCI Insight [Internet]. 2025 Nov 11; Available from: 10.1172/jci.insight.197711

21. Reeves SR, Kolstad T, Lien TY, Herrington-Shaner S, Debley JS. Fibroblast-myofibroblast transition is differentially regulated by bronchial epithelial cells from asthmatic children. Respir Res. 2015 Feb 13;16(1):21.

22. Gruenert DC, Finkbeiner WE, Widdicombe JH. Culture and transformation of human airway epithelial cells. Am J Physiol. 1995 Mar;268(3 Pt 1):L347–60.

23. Albano GD, Zhao J, Etling EB, Park SY, Hu H, Trudeau JB, et al. IL-13 desensitizes β2adrenergic receptors in human airway epithelial cells through a 15-lipoxygenase/G protein receptor kinase 2 mechanism. J Allergy Clin Immunol. 2015 May;135(5):1144–53.e1–9.

24. Zhao J, Maskrey B, Balzar S, Chibana K, Mustovich A, Hu H, et al. Interleukin-13-induced MUC5AC is regulated by 15-lipoxygenase 1 pathway in human bronchial epithelial cells. Am J Respir Crit Care Med. 2009 May 1;179(9):782–90.

25. Reddel HK, Taylor DR, Bateman ED, Boulet LP, Boushey HA, Busse WW, et al. An official American Thoracic Society/European Respiratory Society statement: asthma control and exacerbations: standardizing endpoints for clinical asthma trials and clinical practice. Am J Respir Crit Care Med. 2009 Jul 1;180(1):59–99.

26. Graham BL, Steenbruggen I, Miller MR, Barjaktarevic IZ, Cooper BG, Hall GL, et al. Standardization of Spirometry 2019 Update. An Official American Thoracic Society and European Respiratory Society Technical Statement. Am J Respir Crit Care Med. 2019 Oct 15;200(8):e70–88.

27. Dweik RA, Boggs PB, Erzurum SC, Irvin CG, Leigh MW, Lundberg JO, et al. An official ATS clinical practice guideline: interpretation of exhaled nitric oxide levels (FENO) for clinical applications. Am J Respir Crit Care Med. 2011 Sep 1;184(5):602–15.

28. Bowerman C, Bhakta NR, Brazzale D, Cooper BR, Cooper J, Gochicoa-Rangel L, et al. A Race-neutral Approach to the Interpretation of Lung Function Measurements. Am J Respir Crit Care Med. 2023 Mar 15;207(6):768–74.

29. Lane C, Burgess S, Kicic A, Knight D, Stick S. The use of non-bronchoscopic brushings to study the paediatric airway. Respir Res. 2005 Jun 8;6(1):53.

30. James RG, Reeves SR, Barrow KA, White MP, Glukhova VA, Haghighi C, et al. Deficient Follistatin-like 3 Secretion by Asthmatic Airway Epithelium Impairs Fibroblast Regulation and Fibroblast-to-Myofibroblast Transition. Am J Respir Cell Mol Biol. 2018 Jul;59(1):104– 13.

31. Reeves SR, Kolstad T, Lien TY, Elliott M, Ziegler SF, Wight TN, et al. Asthmatic airway epithelial cells differentially regulate fibroblast expression of extracellular matrix components. J Allergy Clin Immunol. 2014 Sep;134(3):663–70.e1.

32. Lachowicz-Scroggins ME, Boushey HA, Finkbeiner WE, Widdicombe JH. Interleukin-13induced mucous metaplasia increases susceptibility of human airway epithelium to rhinovirus infection. Am J Respir Cell Mol Biol. 2010 Dec;43(6):652–61.

33. Jakiela B, Gielicz A, Plutecka H, Hubalewska-Mazgaj M, Mastalerz L, Bochenek G, et al. Th2-type cytokine-induced mucus metaplasia decreases susceptibility of human bronchial epithelium to rhinovirus infection. Am J Respir Cell Mol Biol. 2014 Aug;51(2):229–41.

34. Doni Jayavelu N, Altman MC, Benson B, Dufort MJ, Vanderwall ER, Rich LM, et al. Type 2 inflammation reduces SARS-CoV-2 replication in the airway epithelium in allergic asthma through functional alteration of ciliated epithelial cells. J Allergy Clin Immunol. 2023 Jul;152(1):56–67.

35. Murphy RC, Lai Y, Altman MC, Barrow KA, Dill-McFarland KA, Liu M, et al. Rhinovirus infection of the airway epithelium enhances mast cell immune responses via epithelial-derived interferons. J Allergy Clin Immunol. 2023 Jun;151(6):1484–93.

36. Vanderwall ER, Barrow KA, Rich LM, Read DF, Trapnell C, Okoloko O, et al. Airway epithelial interferon response to SARS-CoV-2 is inferior to rhinovirus and heterologous rhinovirus infection suppresses SARS-CoV-2 replication. Sci Rep. 2022 Apr 28;12(1):6972.

37. Bhakta NR, Solberg OD, Nguyen CP, Nguyen CN, Arron JR, Fahy JV, et al. A qPCR-based metric of Th2 airway inflammation in asthma. Clin Transl Allergy. 2013 Jul 17;3(1):24.

38. Zhu Z, Homer RJ, Wang Z, Chen Q, Geba GP, Wang J, et al. Pulmonary expression of interleukin-13 causes inflammation, mucus hypersecretion, subepithelial fibrosis, physiologic abnormalities, and eotaxin production. J Clin Invest. 1999 Mar;103(6):779–88.

39. Ravi A, Koster J, Dijkhuis A, Bal SM, Sabogal Piñeros YS, Bonta PI, et al. Interferon-induced epithelial response to rhinovirus 16 in asthma relates to inflammation and FEV. J Allergy Clin Immunol. 2019 Jan;143(1):442–7.e10.

40. Moui A, Klein M, Hassoun D, Dijoux E, Cheminant MA, Magnan A, et al. The IL-15 / sIL15Rα complex modulates immunity without effect on asthma features in mouse. Respir Res. 2020 Jan 29;21(1):33.

41. Verbist KC, Klonowski KD. Functions of IL-15 in anti-viral immunity: multiplicity and variety. Cytokine. 2012 Sep;59(3):467–78.

42. Ramonell RP, Oriss TB, McCreary-Partyka JC, Kale SL, Brandon NR, Ross MA, et al. CD8+ TEMRAs in severe asthma associate with asthma symptom duration and escape proliferation arrest. JCI Insight [Internet]. 2025 Apr 22;10(8). Available from: 10.1172/jci.insight.185061

43. Brightling C, Berry M, Amrani Y. Targeting TNF-alpha: a novel therapeutic approach for asthma. J Allergy Clin Immunol. 2008 Jan;121(1):5–10; quiz 11–2.

44. Berry MA, Hargadon B, Shelley M, Parker D, Shaw DE, Green RH, et al. Evidence of a role of tumor necrosis factor alpha in refractory asthma. N Engl J Med. 2006 Feb 16;354(7):697–708.

45. Choi IW, Sun-Kim, Kim YS, Ko HM, Im SY, Kim JH, et al. TNF-alpha induces the late-phase airway hyperresponsiveness and airway inflammation through cytosolic phospholipase A(2) activation. J Allergy Clin Immunol. 2005 Sep;116(3):537–43.

46. Ray A, Camiolo M, Fitzpatrick A, Gauthier M, Wenzel SE. Are We Meeting the Promise of Endotypes and Precision Medicine in Asthma? Physiol Rev. 2020 Jul 1;100(3):983–1017.

47. Le Floc’h A, Allinne J, Nagashima K, Scott G, Birchard D, Asrat S, et al. Dual blockade of IL-4 and IL-13 with dupilumab, an IL-4Rα antibody, is required to broadly inhibit type 2 inflammation. Allergy. 2020 May;75(5):1188–204.

48. Pesic J, Nieto-Fontarigo JJ, Pardali K, Delaney S, Olsson H, Uller L. T2 cytokine-driven alarmin and antiviral responses in asthma: insights into immune modulation and the role of IL-4Rα targeting. Front Allergy. 2025 Apr 30;6:1576816.

